# Loss of CPAP expression promotes sustained EGFR signaling and Epithelial-Mesenchymal Transition in oral cancer cells

**DOI:** 10.1101/2020.08.05.237107

**Authors:** Radhika Gudi, Harinarayanan Janakiraman, Phillip Howe, Viswanathan Palanisamy, Chenthamarakshan Vasu

## Abstract

Oral squamous cell carcinoma (OSCC) is the most common type of head and neck squamous cell carcinoma (HNSCC). Altered epidermal growth factor receptor (EGFR) levels can contribute to tumor metastasis and resistance to therapies. The epithelial-mesenchymal transition (EMT), by which epithelial cells acquire a mesenchymal and invasive phenotype, contributes significantly to tumor metastasis in OSCC, and EGFR signaling is known to promote this process. Microtubule inhibition therapies cause EGFR inactivation or increase the sensitivity to EGFR targeting drugs in various cancers including OSCC. In this study, using OSCC model, we show that loss of a microtubule/tubulin binding protein, centrosomal protein 4.1-associated protein (CPAP), which is critical for centriole biogenesis and normal functioning of centrosome, caused an increase in the EGFR levels and signaling and, enhanced the EMT features and invasiveness of OSCC cells. Further, depletion of CPAP increased the tumorigenicity of these cells in a xeno-transplant model. Importantly, CPAP loss-associated EMT features and invasiveness of multiple OSCC cells were attenuated upon depletion of EGFR in them. Overall, our novel observations suggest that in addition to its previously known regulatory role in centrosome biogenesis and function, CPAP plays an important role in suppressing EMT and tumorigenesis in OSCC by regulating EGFR homeostasis and signaling.

## Introduction

Head and neck squamous cell carcinoma (HNSCC) represents the sixth most common cancer with more than 600,000 new patients diagnosed worldwide and it is linked to more than 300,000 deaths every year^1^. Most of the head and neck cancers are squamous cell carcinomas (HNSCC) that arise from mucosal surfaces of the oral cavity (OSCC). In most cancers including OSCC, altered epidermal growth factor receptor (EGFR/ErbB1/HER1) levels contribute to tumorigenesis, metastasis, and resistance to therapies and poor rate of patient survival^2-6^. The epithelial-mesenchymal transition (EMT), by which epithelial cells acquire a mesenchymal and invasive phenotype, contributes significantly to these features^7, 8^. EGFR is significantly altered in OSCC and its prolonged signaling is mitogenic, driving the uncontrolled proliferation of tumor cells^9,10^. In addition, EGFR expression and its signal transduction pathways play an important role in determining the sensitivity to chemo or radiotherapy^11, 12^. Importantly, excessive signaling through this receptor also triggers EMT in tumor cells, which makes them more invasive and metastatic, and resistant to chemotherapy^13-16^. Hence, this receptor has become one of the major targets for new therapies being investigated in OSCC^13, 17, 18^. Despite these advances in the understanding of EGFR signaling, the regulatory mechanisms underlying EGFR signaling and their effects on cancer initiation, progression and metastasis are not fully understood.

Recent studies have shown that microtubule inhibition causes EGFR inactivation or increases the sensitivity to EGFR targeting drugs in various cancers including OSCC^19-22^. Microtubules and microtubule-organizing center (MTOC) have multiple roles in cellular functions including homeostasis of cell signaling, formation of cilia, cytoskeletal actin organization, and centrosome/centriole duplication and normal cell division^23^. The centrosome, which contains two centrioles, functions as the major MTOC^24, 25^. Normal centriole duplication is critical for various cellular processes^26-29^. Abnormalities in this process can lead to cellular aging, cancer, decline in stem cell function, ciliopathies, and neuropathies including microcephaly^28, 30-36^. Deregulation in the number of centrioles and centrosomes occurs in cancer and was shown to contribute to tumorigenesis^37-40^. Reports by us^41-43^ and others^28, 44-48^ have shown a pivotal role for key centriole proteins including the microcephaly-associated protein CPAP (SAS-4/LIP1; encoded by CENPJ gene) in centriole duplication. These studies have shown that CPAP, a microtubule binding protein, is indispensable in regulating the centriole and cilia dimensions^28, 41-45, 49-51^. Both overexpression as well as depletion of CPAP causes the generation of multinucleated cells, mitotic errors, abnormalities in bipolar spindle formation, spindle positioning and orientation for normal and asymmetric cell division^26, 52-54^, which are some of the key features of cancer malignancies. Mutations of CPAP have been linked to the aforementioned with centrosome-defect associated disorders^26, 54-61^. It has also been shown that inhibition of CPAP-tubulin/microtubule interaction prevents the proliferation of centrosome‐amplified cancer cells^62^. Nevertheless, if CPAP has a tumor suppressive regulatory role through EGFR levels and signaling dynamics in cancer cells is unknown.

In this study, we observed that CPAP depletion led to elevated cellular levels of, and signaling by, EGFR in them. Interestingly, CPAP-loss alone was sufficient to cause spontaneous EMT-like features in OSCC cells. In addition, CPAP-depleted OSCC cells showed increased invasiveness and tumorigenicity, proving the tumor suppressor role of CPAP. EGFR depletion suppressed the EMT-features in as well as the invasiveness of CPAP-depleted OSCC cells suggesting that the CPAP-EGFR axis contributes to the EMT phenotype. Paradoxically, however, we show that EGFR signaling, which is known to promote EMT^63-65^, upregulates the cellular levels of CPAP in OSCC cells. Our studies reveal a new mechanism of tumor suppression by the centriole-associated protein, CPAP. Overall, this study suggests that while higher centrosome amplification and aberrant centriole features are frequently reported phenotypes in cancer^62, 66-68^ is possibly the consequence of tumor progression-associated inflammation, CPAP exerts tumor preventive function in OSCC through promoting the homeostasis of EGFR and suppressing EMT.

## Materials and Methods

### Cell culture

Oral cancer cell lines were acquired from ATCC or obtained from the laboratory of Dr. Thomas Carey, University of Michigan. Immortalized human normal oral keratinocytes OKF6tert1 (OKF6) cell line was obtained from Dr. Jim Rheinwald, Harvard Medical School. Most studies were focused on SCC-Cal27 (adenosquamous carcinoma cell line) and UM-SCC-74A and UM-SCC-74B which are mesenchymal in origin and they were cultured in Dulbecco’s modified Eagle medium that was supplemented with 10% fetal bovine serum and other additives such glutamine, sodium pyruvate, bicarbonate, minimum essential amino acids and antibiotics/antimycotic. Intracellular CPAP levels were also tested in other oral cancer cell lines such as UM-SCC1A, UM-SCC9, UM-SCC4, UM-SCC25, UM-SCC11A, some of which were cultured in DMEM/F12 medium containing hydrocortisone and above-mentioned supplements. Cells were cultured in a 37°C incubator with 5% CO2.

### Constructs, transfection and generation of stable cell lines

CPAP-depleted stable oral cancer cells were generated using the pLKO.1-TRC-puromycin lentiviral vector based system. Constitutive and doxycycline inducible vectors expressing CPAP-specific and scrambled control-shRNAs, showing validated 80% CPAP knockdown were purchased from the Sigma Mission shRNA library. EGFR specific or scrambled control siRNAs were purchased from Santa Cruz Biotech. Doxycycline was purchased from Fisher. Transfections were performed using the TransIT-X2 reagent purchased from MirusBio.

### Antibodies

Commercial antibodies were purchased from: CPAP/CENPJ, Actin, Zeb, Slug, N-cadherin and E-cadherin (Proteintech), vimentin (Cell Signaling Technology), and EGFR (Santa Cruz Biotech). Secondary HRP-linked anti-mouse and -rabbit antibodies were purchased from Biorad and Amersham respectively and Alexa-conjugated antibodies were from Invitrogen.

### Cell lysis and Western blotting

Whole cell lysates were prepared using the RIPA lysis buffer containing protease inhibitors, centrifuged, and supernatants were used for further analysis. Lysates were then subjected to immunoblotting after separation by SDS-PAGE polyacrylamide gel electrophoresis and Western blotting transfer onto PVDF membrane (Biorad), followed by immunoblotting with standard techniques.

### Immunofluorescence

Cells grown on coverslips were fixed with 4% paraformaldehyde and permeabilized using 0.1% saponin containing buffer for 30 mins. Blocking with 1% BSA as well as primary and secondary antibody dilutions were made in permeabilization buffer and incubations were done at 37°C. Images were acquired using the Zeiss 880 confocal microscope using the 63X oil immersion objective with n.a. 1.4.

### Migration and invasion assay

Oral cancer cells treated differently were trypsinized, counted and an equal number were seeded in serum free media in the upper transwell chamber with or without matrigel (filter pore size 8μm, BD). 5% FBS-containing media was loaded in the bottom chamber as chemoattractant. After 24h incubation at 37°C, cells on opposite side of filter were fixed with 2% glutaraldehyde. Cells on the seeded membrane side were scraped off using a wet cotton swab, followed by staining of the membrane with 2% crystal violet stain. After numerous washes, filters were dried, membranes were cut and mounted on glass slides in immersion oil. Number of cells in various fields were imaged using 10X Omax objective and imaged using the Toupview software. Average number of cells in various fields between 3 independent experiments has been plotted.

### Tumor microarray analysis (TMA) patient tissues

Human tumor and adjacent normal tissues were collected from the Biorepository & Tissue Analysis at Hollings Cancer Centre and stained and scored blindly and imaged by the IHC core.

### Tumor xenografts

Two million, doxycycline inducible Cal27/74B-shControl and -shCPAP were suspended in Matrigel to make it 50% concentration and injected subcutaneously in both the right and left flanks of 6-week-old male and female athymic nude mice. After sacrifice, tumors were removed and weighed. All animal experiments were approved by the Medical University of South Carolina Institutional Animal Care and Use Committee (Charleston, SC).

## Results

### CPAP depletion endows OSCC cells with EMT phenotype and properties

While conducting centriole biogenesis-related studies using HeLa cells, we observed that depletion, but not overexpression, of CPAP resulted in a typical EMT-like morphological change (not shown). Since OSCC is a highly prevalent cancer, we examined EMT features in OSCC cells upon CPAP depletion. Stable CPAP depletion was performed in 3 oral cancer cell-lines with: 1) an epithelial phenotype (SCC-Cal27), 2) a mesenchymal phenotype of primary tumor origin (UM-SCC-74A), and 3) a mesenchymal phenotype of recurrent tumor origin (UM-SCC-74B). Stable CPAP-depleted and control cell-lines were generated by transduction using pLKO.1-puro lentiviral vectors encoding CPAP shRNA and scrambled control shRNA respectively, followed by selection using puromycin. As observed in Fig. **1A**, all three oral cancer cell-lines, with epithelial and mesenchymal phenotypes that were transduced with CPAP shRNA showed elongated morphology, a key feature of EMT, compared to control shRNA expressing cells. The spindle-like stretched appearance was more prominent in the CPAP-devoid mesenchymal cells. Immunoblot (IB) analysis of these cells revealed the elevated expression of one or more mesenchymal markers (vimentin, N-cadherin, Zeb and Slug) in all three cell types upon CPAP depletion compared to their control counterparts (Fig. **1B**). Although SCC-Cal27 did not show detectable levels of vimentin, diminished levels of epithelial marker E-cadherin and higher expression of transcription factors Zeb and Slug were detected in these cells upon CPAP depletion. On the other hand, both UM-SCC-74A and UM-SCC-74B cells with CPAP depletion showed higher levels of mesenchymal markers vimentin, N-cadherin, Zeb1 and Slug. Expression levels of E-cadherin and vimentin were also examined by immunofluorescence microscopy. As observed in **Fig. 1C**, while CPAP-depleted SCC-Cal27 cells expressed relatively lower levels of E-cadherin, CPAP-depleted UM-SCC-74A and UM-SCC-74B cells expressed higher levels of vimentin compared to respective control cells. These observations suggest that loss of CPAP expression renders cells more susceptible to undergo EMT.

**Fig. 1.**
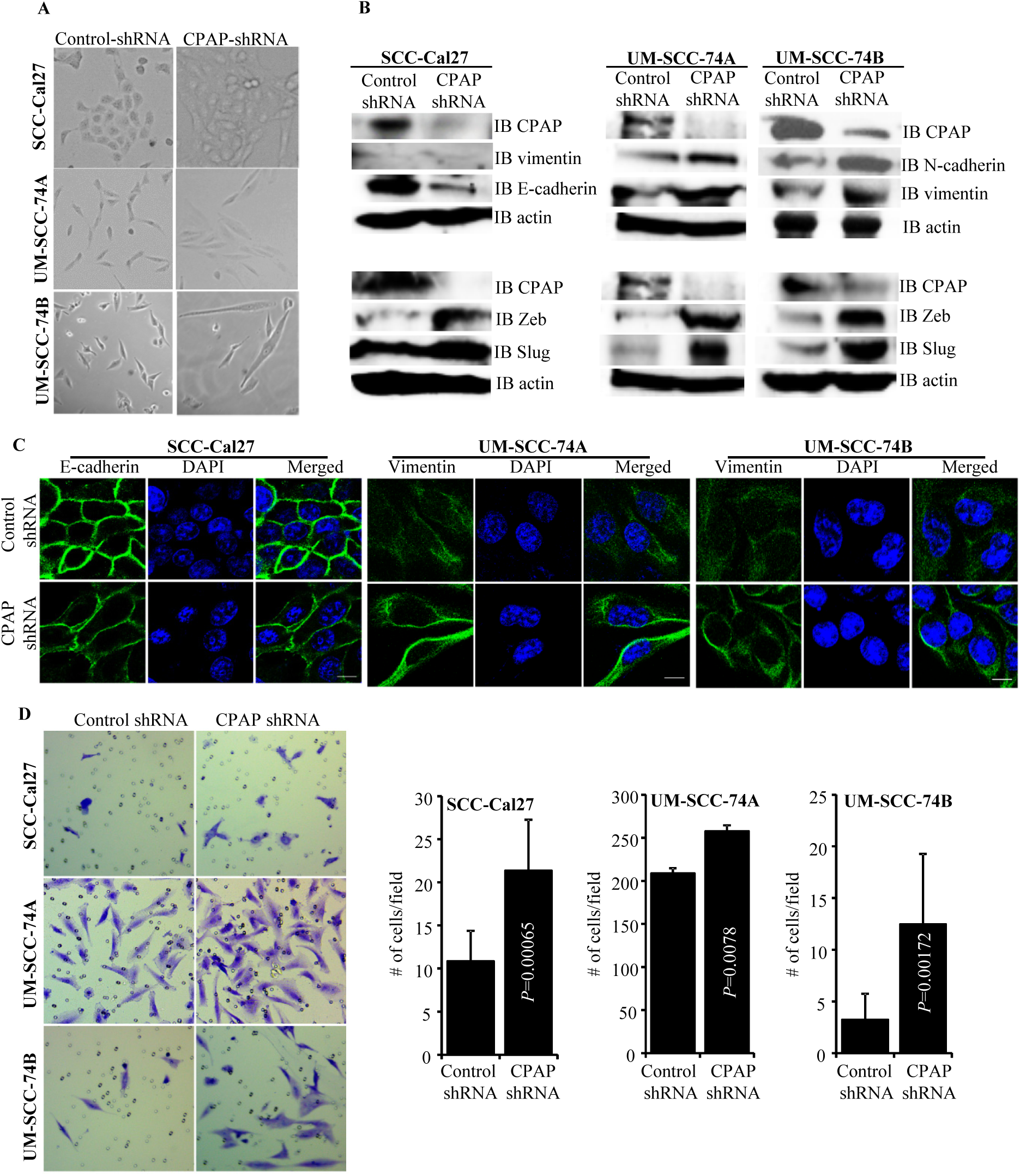
CPAP depleted cells show spontaneous EMT-like morphology and upregulated mesenchymal protein expression. **A)** Representative bright-field images of indicated oral cancer cell lines transduced with control-shRNA or CPAP-shRNA expressing lentiviral particles and selected using puromycin (for a week) for stable shRNA expression. **B)** IB showing protein levels of various EMT associated markers along with CPAP and β-actin in indicated cell lines that are stably expressing control-shRNA and CPAP-shRNA. **C)** Control-shRNA and CPAP-shRNA expressing OSCC cells were fixed and permeabilized, and subjected to immunofluorescence microscopy to detect E-cadherin or vimentin (green) and nuclear stain DAPI (blue). Scale bar: 10μm. **D)** Transwell-membrane plates with matrigel coating were used to determine the invasive properties of OSCC cell-lines. Equal numbers of control-shRNA and CPAP-shRNA expressing indicated cell-lines were seeded in serum free media in the upper chamber and incubated for 24h. Cells on lower chamber side of the transwell membrane were stained with crystal violet, imaged and the average number (mean±SD) of cells from multiple fields were plotted. Representative results from one of the three independent experiments are shown. *P*-value by Mann-Whitney test.

Since, cells undergoing the EMT process are known to be more invasive, control-shRNA and CPAP-shRNA expressing cells were subjected to cell migration and invasion assay using trans-well inserts with and without matrigel coating. While the migratory properties of control and CPAP-depleted cells in membrane-well plates without matrigel coating were not different (not shown), all three OSCC cell lines with CPAP depletion showed significantly higher matrigel invasion ability compared to control cells **(Fig. 1D)**. Collectively, these observations suggest that CPAP plays a role in inhibiting the EMT process in OSCC cells.

### CPAP loss enhances OSCC cell-line induced tumor growth in vivo

Further, to test if the EMT-like features and enhanced invasiveness of CPAP-depleted OSCC cells translate into rapid tumor growth in vivo, SCC-Cal27, UM-SCC-74A and UM-SCC-74B cells stably expressing control-shRNA and CPAP-shRNA under a doxycycline inducible promoter **(Fig. 1A)** were injected s.c. into an athymic-nude mice. These mice were given doxycycline in drinking water and monitored for tumor growth at timely intervals. Tumor growth was relatively rapid (not shown) in CPAP-shRNA expressing, both SCC-Cal27 (epithelial) and UM-SCC-74B (mesenchymal), cell recipient groups compared to their control-shRNA recipient counterparts. Importantly, the average tumor weights upon euthanasia of SCC-Cal27 cell recipients on day 48 and UM-SCC-74B cell recipients on day 24 post-injection were significantly higher in CPAP-shRNA expressing cell recipients compared to respective controls **(Fig. 1B)**. Of note, in our hands, in spite of the mesenchymal phenotype, UM-SCC-74A cells failed to induce tumor in mice even after 50 days post-injection (not shown). In a parallel experiment, a small cohort of mice received s.c. injection of UM-SCC-74B cells that overexpress CPAP under doxycycline inducible promoter and were monitored similarly. Tumor weights in mice that received CPAP overexpressing cells were relatively lower compared to control cell recipients **(Supplemental Fig. 1)**. Overall, these results along with our in vitro findings demonstrating that CPAP knockdown enhances EMT properties and invasiveness (Fig. 3) show that CPAP suppresses the tumorigenic properties of OSCC cells.

**Fig. 3:**
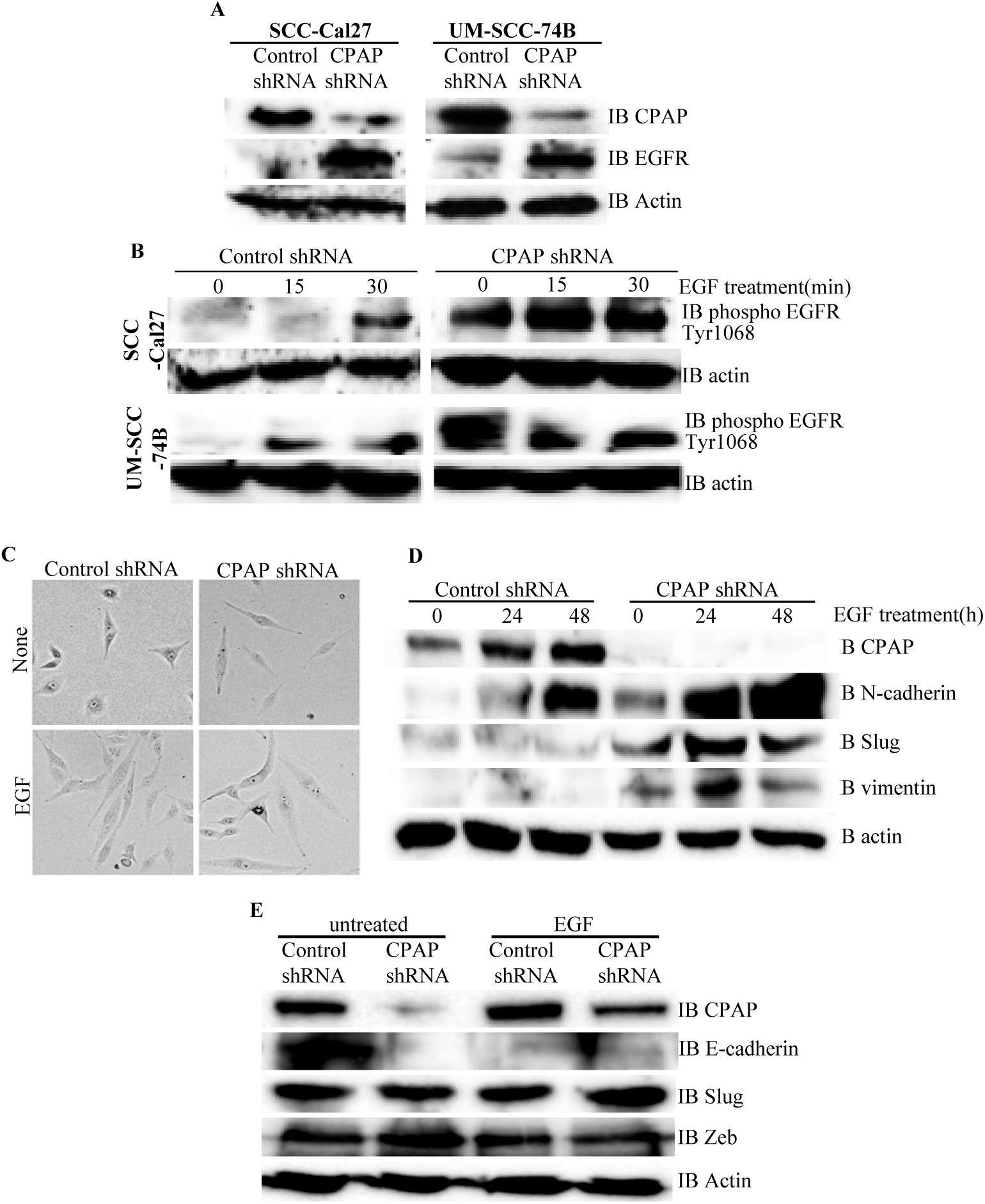
CPAP depletion causes higher cellular levels of total and phospho-EGFR proteins and EGF treatment enhances EMT-like features in CPAP-depleted OSCC cells. **A)** Indicated OSCC cell lines that are stably expressing control shRNA or CPAP shRNA were subjected to IB to detect total EGFR protein and β-actin. **B)** These control and CPAP-depleted cell lines were maintained in serum-free media overnight, treated with cycloheximide for 1h and treated with EGF (30ng/ml), and incubated at 37°C for indicated durations to induce signaling. Levels of phosphorylated EGFR (Tyr1068) were detected in protein equalized cell lysates by IB. **C)** Representative bright-field microscopy images of control and CPAP depleted UM-SCC-74B cells left untreated or treated with EGF (30 ng/ml) for 48h are shown. **D)** IB analysis of control or CPAP-depleted UM-SCC-74B that were left untreated or treated with EGF for indicated time points and subjected to IB to detect the levels of EMT-associated proteins N-cadherin, vimentin and Slug. E-cadherin levels were undetectable in these cells. **E)** Control and CPAP-depleted SCC-Cal27 cells were also subjected to EGF treatment for 48 h and subjected to IB to detect E-cadherin, Zeb and Slug. Vimentin and N-Cadherin were undetectable in these cells.

### CPAP loss enhances the total and phospho-EGFR levels in OSCC cells

Since EGF treatment induced EGFR signaling triggers EMT features in OSCC cells^63, 69^, EGFR levels in CPAP depleted cells were examined. As observed in **Fig. 3A**, CPAP depletion resulted in a profound increase in the cellular EGFR protein levels in all the three tested OSCC cell-lines (SCC-Cal27, UM-SCC-74B and UM-SCC-74B) with either epithelial or mesenchymal phenotypes. Next, we treated these cells with EGF and examined for the cellular levels of phosphorylated EGFR levels, in protein equalized cell lysates, as an indication of active signaling by this receptor. Fig. **3B** shows that, as expected, EGF treatment resulted in an increase in the levels of phosphorylated EGFR in control SCC-Cal27, UM-SCC-74B and UM-SCC-74B cells. Interestingly, the basal levels of phospho-EGFR were profoundly higher in CPAP-depleted cells to begin with, compared to control cells. Further, the increase in phospho-EGFR levels upon EGF treatment appeared to be more rapid and striking in CPAP-depleted cells, particularly in SCC-Cal27 and UM-SCC-74A cells, compared to their control counterparts. Collectively, these observations suggest that CPAP has a crucial role in maintaining the homeostasis of growth factor receptors like EGFR and loss of CPAP results in, potentially, enhanced and persistent signaling through this receptor, causing EMT and enhancing the tumorigenic properties of OSCC cells.

### EGFR activation under CPAP loss enhances EMT features of OSCC cells

Since phospho-EGFR levels were higher in CPAP depleted OSCC cells and EGF treatment of these cells caused a rapid increase in this phospho-protein levels, the EMT features of untreated and EGF treated cells were examined. As observed in **Fig. 3C**, UM-SCC-74B cells with and without CPAP knockdown showed the classic spindle-like elongated EMT-associated morphology upon EGF treatment compared to respective control cells. While the untreated CPAP-depleted cells showed spontaneous EMT features (as mentioned in Fig. 2A), these morphologic changes were more pronounced upon EGF treatment. Examination of EMT associated markers in these cells revealed a profound upregulation of N-cadherin, but not Slug and Vimentin, levels in control cells upon EGF treatment. However, CPAP depleted cells showed, in addition to higher basal levels compared to control cells, an increase in the levels of N-cadherin, Slug and Vimentin upon EGFR treatment **(Fig. 3D)**. Similarly, although the classic EMT associated elongated appearance was not observed with control epithelial SCC-Cal27 cells upon EGF treatment alone (not shown), E-cadherin levels were diminished in these control cells upon EGF treatment. Importantly, upon EGF treatment, an increase in the protein levels of Slug was observed only in CPAP depleted, but not control, SCC-Cal27 cells **(Fig. 3E)**. These observations support the notion that enhanced EGFR signaling is responsible for the EMT-like features of CPAP depleted OSCC cells.

**Fig. 2.**
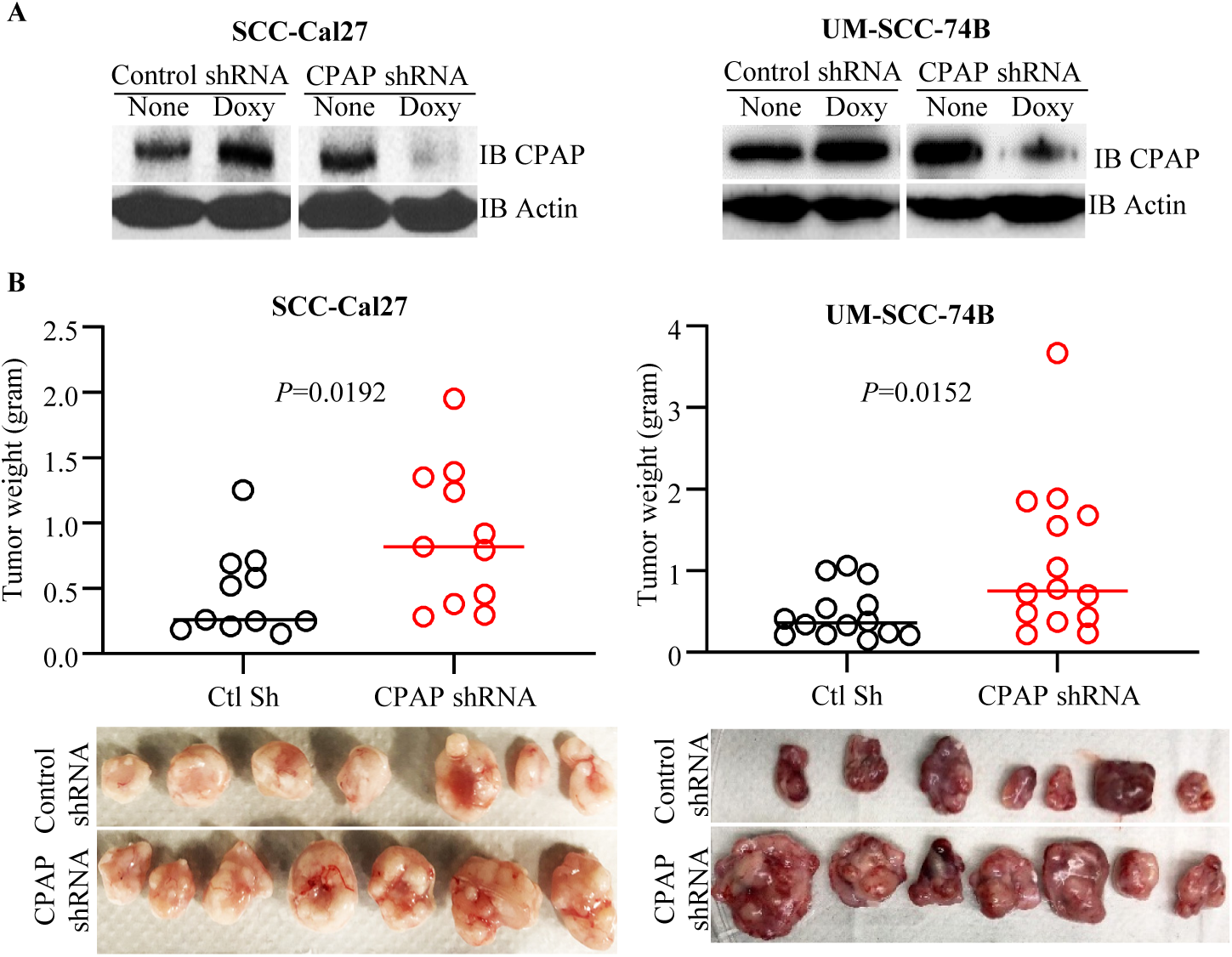
CPAP depletion enhances the tumorigenic properties of OSCC cell lines. SCC-Cal27 and UM-SCC-74B cell lines were transduced with control-shRNA or CPAP-shRNA (under doxy-inducible promoter) lentiviral particles, and the cells with stable integration were selected using puromycin. **A)** shRNA expression was induced in these cells by culturing for 72h in the presence of doxy (10 μg/ml) and subjected to IB to detect CPAP and β-actin. **B)** Eight week old nude mice were injected s.c. with these control-shRNA and CPAP-shRNA lentivirus transduced cells (approx: 2×10^6^ cell/mouse), mixed with equal volume of matrigel on the flank, monitored for tumor progression, and euthanized after 24 days (UM-SCC-74B cell recipients) or 48 days (SCC-Cal27 cell recipients) to determine the tumor weight. Cumulative data from two independent experiments (total of 11 mice/group for SCC-Cal27 cells and 14 mice/group for UM-SCC-74B cell recipients) are shown (B; upper panels). Images of tumors harvested from one experiment are also shown (B; lower panels). *P*-value by Mann-Whitney test.

### Depletion of EGFR results in elimination of CPAP-loss induced EMT features and invasiveness of OSCC cells

To assess the role of EGFR signaling in CPAP-depletion associated EMT features, control and CPAP depleted SCC-Cal27, UM-SCC-74A and UM-SCC-74B cells were treated with scrambled control or EGFR specific siRNA. As shown in **Fig. 4A**, EGFR depletion in control SCC-Cal27 and UM-SCC-74A cells did not affect, or caused only a modest increase in, the expression levels of transcription factors Slug and Zeb compared to control siRNA treated cells. In control UM-SCC-74B cells, EGFR depletion caused the suppression of protein levels of Slug, but not Zeb. Interestingly, under CPAP deficiency, EGFR depletion caused the suppression of Zeb and Slug levels in SCC-Cal27 cells, Zeb in UM-SCC-74A cells and Slug in UM-SCC-74B cells. We then examined if these differential changes in EMT marker levels upon EGFR depletion in CPAP depleted cells impacts their migratory and invasive properties by plating equal number of specific cell types in trans-wells without and with matrigel coating. OSCC cell lines with both epithelial phenotype (SCC-Cal27) and mesenchymal phenotype (UM-SCC-74B and UM-SCC-74B) showed comparable migratory properties, based on the assay using trans-wells without matrigel, irrespective of CPAP and/or EGFR deficiency **(Fig. 4B)**. However, while CPAP deficiency caused enhanced invasion by all three cell-lines in a matrigel transwell assay as per our observations of Fig.1, CPAP-deficient SCC-Cal27, UM-SCC-74B and UM-SCC-74B cells with EGFR depletion showed profoundly suppressed invasiveness compared to their counterparts without EGFR depletion **(Fig. 4C)**. Overall, these observations confirm that CPAP loss associated EMT phenotype and enhanced tumorigenic properties of OSCC cells are EGFR-, and its enhanced signaling, -dependent.

**Fig. 4.**
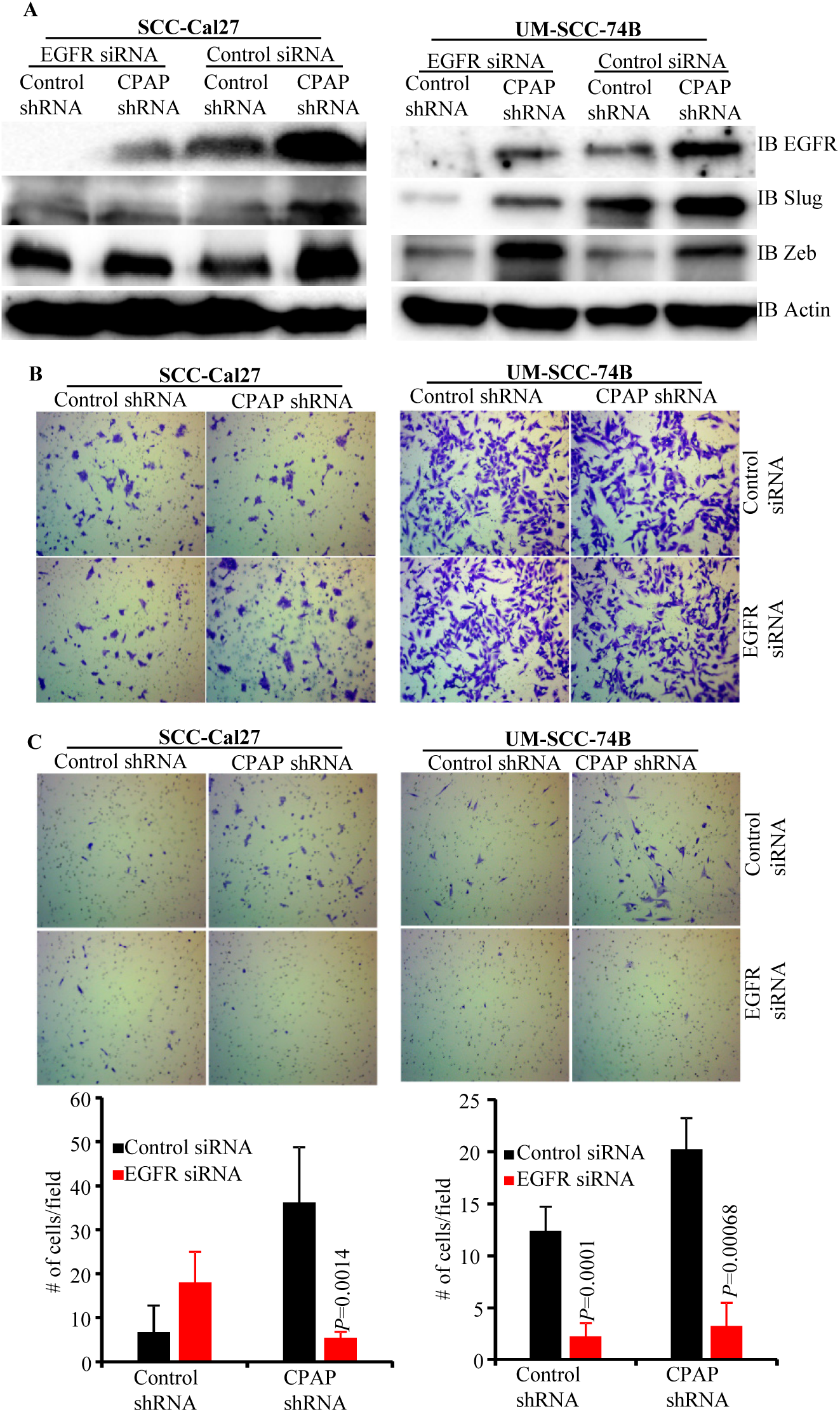
EGFR depletion diminishes the EMT phenotype and invasive property of CPAP-depleted OSCC cells. Control-shRNA and CPAP-shRNA expressing indicated cells were treated with control-siRNA or EGFR-siRNA for 48h. A) Cells were subjected to IB to detect EMT associated transcription factors Slug and Zeb along with EGFR and β-actin. Equal number of each cell types were also subjected to migration and matrigel-invasion assay using the transwell approach and imaged. **B)** Representative fields of migrated cells on insert membranes with no matrigel coating are shown. **C)** Representative fields of invasive cells on membranes with matrigel coating (upper panels) and average number (mean±SD) of cells of multiple fields (lower panels) are shown. Representative results from one of the three independent experiments are shown. *P*-value by Mann-Whitney test.

### CPAP protein levels in OSCC cell-lines under EMT inducing conditions and steady state

To determine if there is a correlation between CPAP protein levels and the phenotype of OSCC cells, we examined the expression levels of CPAP and EMT associated markers in OSCC cell-lines and immortalized, normal human oral keratinocyte (OKF6). As observed in **Fig. 5A**, when protein equalized cell lysates were employed, UM-SCC-1A and SCC-Cal27 along with normal OKF6 cells showed relatively higher expression of epithelial marker E-cadherin. However, UM-SCC-74A, UM-SCC-74B and UM-SCC-9 expressed higher levels of mesenchymal markers vimentin and N-cadherin. While UM-SCC-25 cells expressed higher levels of both epithelial and mesenchymal markers, UM-SCC-11A expressed these markers at very low levels. Importantly, however, CPAP protein levels did not show a clear correlation with the EMT marker levels. To determine the impact of growth factors that are known to promote EMT and tumorigenesis^69-73^ on cellular levels of CPAP, OKF6 cells and oral cancer cells (with epithelial and mesenchymal phenotypes, UM-SCC-Cal27 and UM-SCC-74B respectively) were treated with EGF and TGFβ1. As observed in **Fig. 5B**, all three cell lines expressed higher levels of CPAP in both EGF and TGFβ1 treated cells, within 48h of exposure. Further, as anticipated EGF and TGFβ1 treatments not only decreased the levels of epithelial marker E-cadherin in OKF6 cells, but also caused an increase in the levels of mesenchymal marker N-cadherin in UM-SCC-74B **(Fig. 5C)**. These observations suggest that while CPAP protein levels could be dependent on cell-lines and their various other properties, EMT inducing conditions may contribute to the cellular accumulation of this protein.

**Fig. 5.**
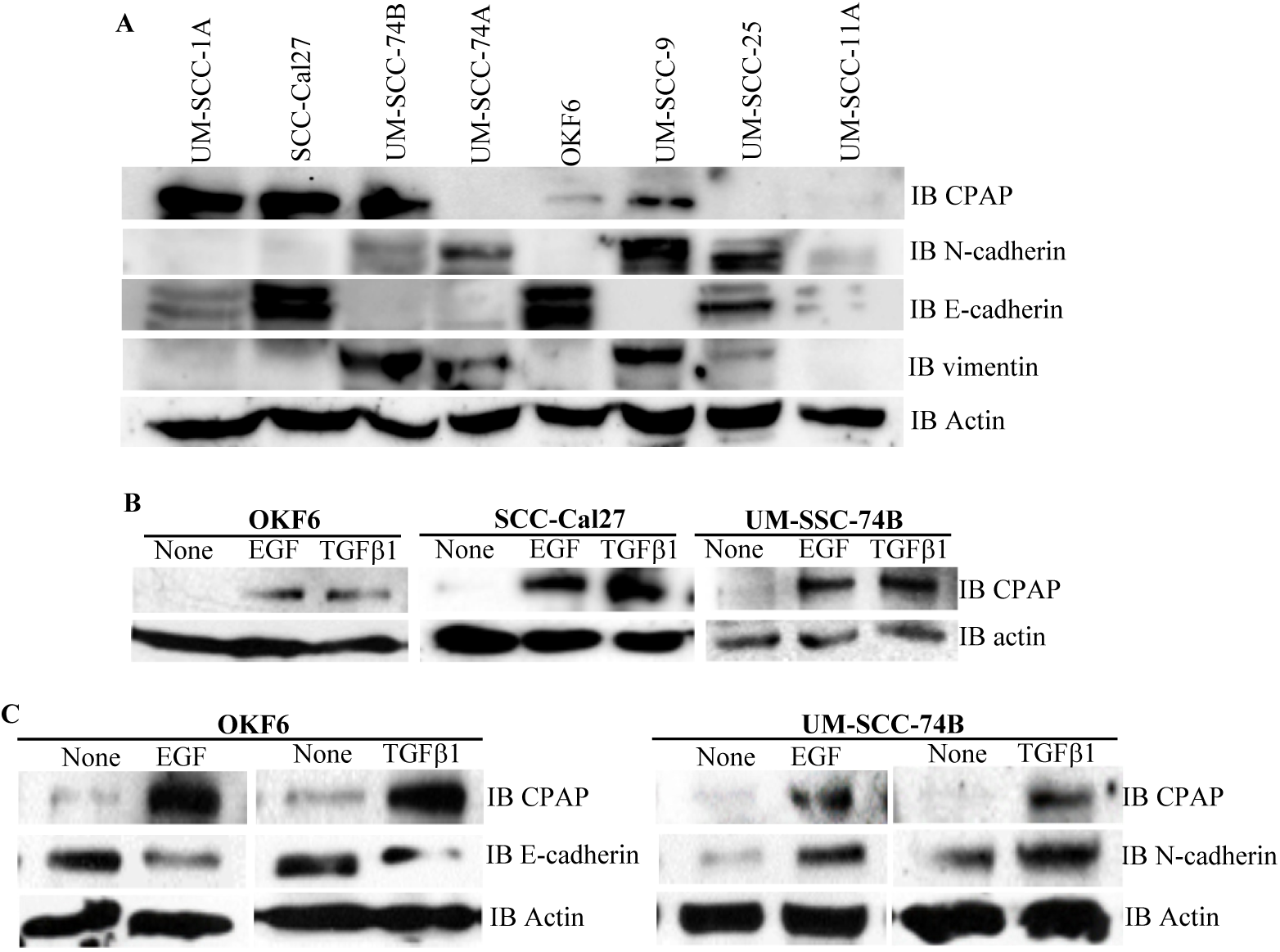
EMT inducing factors EGF and TGF treatments upregulate CPAP protein levels in OSCC cell lines. **A)** IB showing protein levels of CPAP, the EMT markers N-cadherin, E-cadherin and vimentin, and β-actin in protein equalized lysates of various OSCC cell-lines. **B)** IB showing CPAP and β-actin protein levels in the indicated OSCC cell lines upon treatment with EGF (30ng/ml) and TGFβ1 (5 ng/ml) for 48h. **C)** IB showing EMT marker E-cadherin or N-cadherin level, along with CPAP and β-actin levels, in OKF6 normal and UM-SCC-74B OSCC cell lines.

## Discussion

Here, using the OSCC model, we demonstrate a negative regulatory role for CPAP, an essential centriole biogenesis protein^54^, in tumor prevention by keeping the EGFR signaling and EMT at bay. CPAP is a microtubule and α-tubulin binding protein and the tubulin-binding property ^49, 74, 75^ is important for its function on the centrioles, especially in restricting the centriole length^28, 45, 52^. In addition, CPAP is required for spindle orientation, which defines normal and asymmetric cell divisions, abnormalities, which could lead to various clinical conditions including tumor malignancies^26, 52-54^. Recently, it has been shown that inhibition of CPAP-tubulin interaction prevents proliferation of centrosome‐amplified cancer cells^62^. However, a role for CPAP as a tumor suppressor in regulating EGFR homeostasis and signaling, and EMT is not known. In this report, we show that although EGFR activation increases the cellular levels of CPAP, loss of CPAP not only increased the total and phosphorylated EGFR levels, but also EGFR dependent-EMT in OSCC. Our results shed light on a novel mechanism of tumor suppression by the centriole-associated protein CPAP.

Centrosome amplification and higher expression of centrosomal proteins are key features of many cancers^62, 66-68^. However, whether these feature are the cause of OSCC and other cancers, or a consequence of tumor progression associated events is largely unknown. Tumor microenvironment is inflammatory and known to express higher levels of growth and cell transformation factors such as TGFβ1 and EGF, and/or their receptors^76-80^. In fact, our results also show that EGF and TGF treatments, which are known to promote the tumorigenic potential of cells, can not only induce EMT-like phenotype, but also increase the expression levels of CPAP in cancer cells. On the other hand, the observation that loss of CPAP results in enhanced EGFR levels and signaling, and EMT features in OSCC cells suggest that this microtubule/tubulin interacting protein negatively regulates the onset of EMT through promoting EGFR homeostasis.

EGFR overexpression was detected in majority of OSCC tumors and many other malignancies^81, 82^. More than 90% of OSCC overexpress EGFR^83, 84^. Associations have been made between the higher expression levels of EGFR and an aggressive phenotype, poor prognosis and resistance to anticancer therapy of OSCC^81^. A number of EGFR targeted therapies have been developed for the treatment of advanced OSCC^85, 86^. Several studies have demonstrated that EGF can induce EMT, a process by which epithelial cells adopt a mesenchymal phenotype or fibroblast-like properties and increase the invasiveness, in cancer cells, eventually leading to metastasis^63, 69, 72^. In general, cells undergoing EMT exhibit down-regulation of the epithelial marker E-cadherin and up-regulation of mesenchymal markers such as N-cadherin and vimentin and the transcription factors Zeb, Slug, Snail. Further, cancer cells undergoing EMT switch from their epithelial morphology and characteristics such as non-motile and non-invasiveness to their mesenchymal elongated, motile, and invasive characteristics^87^. Our observations show that loss of CPAP in OSCC cells not only increases the EGFR levels and signaling as indicated by phospho-protein levels, but the CPAP-EGFR axis also endows them with EMT like features, enhanced invasiveness and tumorigenesis.

It has been shown that not only does EGF signaling promote EMT in OSCC cells^63, 69^, but also that these cancer cells that are undergoing EMT express considerably lower levels of EGFR and are less susceptible to EGFR targeted therapies, which leads to their chemotherapeutic resistance^88^. Interestingly, while CPAP depletion associated increase in total EGFR appears to be similar in three different cell lines tested, the dynamics of phosphorylated EGFR levels upon ligand activation appear to be different. Basal level of phospho-EGFR upon CPAP depletion was found to be profoundly higher in mesenchymal cell-line UM-SCC-74B which was derived from a recurrent metastatic tumor and in an OSCC cell line SCC-Cal27 with epithelial phenotype, compared to mesenchymal UM-SCC-74A which is of a primary metastatic tumor origin. However, while ligand activation caused the rapid induction of phospho-EGFR in CPAP depleted UM-SCC-74A, its levels appeared to be downregulated rapidly in UM-SCC-74B cells upon ligand activation in vitro. Further, ligand engagement associated increase in phospho-EGFR in CPAP depleted SCC-Cal27 was modest compared to UM-SCC-74A, albeit clearly higher than its respective control. These observations suggest that rapid tumor inducing property of UM-OSCC-74B could be due to low threshold activation and persistent signaling of EGFR in these cells, especially under CPAP deficiency. These dynamics could also explain why these three oral cell lines with different phenotypic and tumor inducing properties express EMT associated markers, Zeb and Slug particularly, differently under CPAP deficiency and upon EGFR depletion. Nevertheless, the common features of CPAP depleted OSCC cells are higher and persistent EGFR expression, enhanced invasiveness and/or tumorigenic ability. Importantly, results from our EGFR depletion studies show that these CPAP-deficiency associated effects are, in fact, EGFR- and, perhaps, its persistent signaling-dependent.

Overall, our observations, reveal that CPAP, a tubulin interacting protein which is critical for centriole biogenesis, has a preventive role in EMT and tumorigenesis by promoting EGFR homeostasis and diminishing the EGFR signaling. While it has been reported before that EGF stimulation endows OSCC cells with stem cell-like properties, increased invasiveness, and tumorigenic properties^63, 69^, the molecular mechanisms underlying the regulation of EGF induced EMT and tumorigenicity were not known. Further, although many studies have shown that microtubule inhibition causes EGFR inactivation or increases the sensitivity to EGFR targeting drugs in various cancers including OSCC^19-22^, whether centrosomes and centriolar proteins contribute to this effect was not investigated before. Hence, this study begins to shed light on the molecular mechanisms by which centrosome/MTOC associated proteins are involved in preventing tumorigenesis.

## Acknowledgements

We would like to thank Dr. T.K. Tang and Dr. Pierre Gonczy for sharing the myc-CPAP/ GFP-CPAP and doxycycline inducible GFP-CPAP expression constructs respectively. We would also like to thank the Cell & Molecular Imaging Shared Resource which is supported by the Hollings Cancer Center, Medical University of South Carolina (P30 CA138313) and the Shared Instrumentation Grant S10 OD018113. This work was supported by NIH grants R21DE026965 and R21DE026965-02S1 to R.G. and C.V. and MUSC internal funds to C.V.

## Data availability

The datasets generated during and/or analyzed during the current study are available from the corresponding author on reasonable request.

## References

1. Vigneswaran, N. & Williams, M.D. Epidemiologic trends in head and neck cancer and aids in diagnosis. Oral Maxillofac Surg Clin North Am 26, 123–141 (2014).

2. Ogiso, H. et al. Crystal structure of the complex of human epidermal growth factor and receptor extracellular domains. Cell 110, 775–787 (2002).

3. Vieira, A.V., Lamaze, C. & Schmid, S.L. Control of EGF receptor signaling by clathrin-mediated endocytosis. Science 274, 2086–2089 (1996).

4. Lafky, J.M., Wilken, J.A., Baron, A.T. & Maihle, N.J. Clinical implications of the ErbB/epidermal growth factor (EGF) receptor family and its ligands in ovarian cancer. Biochim Biophys Acta 1785, 232–265 (2008).

5. Roepstorff, K., Grovdal, L., Grandal, M., Lerdrup, M. & van Deurs, B. Endocytic downregulation of ErbB receptors: mechanisms and relevance in cancer. Histochem Cell Biol 129, 563–578 (2008).

6. Khattri, A. et al. Rare occurrence of EGFRvIII deletion in head and neck squamous cell carcinoma. Oral Oncol 51, 53–58 (2015).

7. Wangmo, C., Charoen, N., Jantharapattana, K., Dechaphunkul, A. & Thongsuksai, P. Epithelial-Mesenchymal Transition Predicts Survival in Oral Squamous Cell Carcinoma. Pathol Oncol Res (2019).

8. Krisanaprakornkit, S. & Iamaroon, A. Epithelial-mesenchymal transition in oral squamous cell carcinoma. ISRN Oncol 2012, 681469 (2012).

9. Tomas, A., Futter, C.E. & Eden, E.R. EGF receptor trafficking: consequences for signaling and cancer. Trends Cell Biol 24, 26–34 (2014).

10. Perez, R., Crombet, T., de Leon, J. & Moreno, E. A view on EGFR-targeted therapies from the oncogene-addiction perspective. Front Pharmacol 4, 53 (2013).

11. Kwok, T.T. & Sutherland, R.M. Differences in EGF related radiosensitisation of human squamous carcinoma cells with high and low numbers of EGF receptors. Br J Cancer 64, 251–254 (1991).

12. Shin, D.M. et al. Epidermal growth factor receptor-targeted therapy with C225 and cisplatin in patients with head and neck cancer. Clin Cancer Res 7, 1204–1213 (2001).

13. Mizumoto, A. et al. Induction of Epithelial-Mesenchymal Transition via Activation of Epidermal Growth Factor Receptor Contributes to Sunitinib Resistance in Human Renal Cell Carcinoma Cell Lines. J Pharmacol Exp Ther 355, 152–158 (2015).

14. Misra, A., Pandey, C., Sze, S.K. & Thanabalu, T. Hypoxia activated EGFR signaling induces epithelial to mesenchymal transition (EMT). PLoS One 7, e49766 (2012).

15. Holz, C. et al. Epithelial-mesenchymal-transition induced by EGFR activation interferes with cell migration and response to irradiation and cetuximab in head and neck cancer cells. Radiother Oncol 101, 158–164 (2011).

16. Talasila, K.M. et al. EGFR wild-type amplification and activation promote invasion and development of glioblastoma independent of angiogenesis. Acta Neuropathol 125, 683–698 (2013).

17. Ito, F. & Takeuchi, K. Novel aspects of epidermal growth factor receptor in relation to tumor development. FEBS J 277, 300 (2010).

18. Lo, H.W. et al. Epidermal growth factor receptor cooperates with signal transducer and activator of transcription 3 to induce epithelial-mesenchymal transition in cancer cells via up-regulation of TWIST gene expression. Cancer Res 67, 9066–9076 (2007).

19. Failly, M. et al. Combination of sublethal concentrations of epidermal growth factor receptor inhibitor and microtubule stabilizer induces apoptosis of glioblastoma cells. Mol Cancer Ther 6, 773–781 (2007).

20. Wu, X. et al. Microtubule inhibition causes epidermal growth factor receptor inactivation in oesophageal cancer cells. Int J Oncol 42, 297–304 (2013).

21. Kitahara, H. et al. Eribulin sensitizes oral squamous cell carcinoma cells to cetuximab via induction of mesenchymal-to-epithelial transition. Oncol Rep 36, 3139–3144 (2016).

22. Chin, T.M. et al. Targeting microtubules sensitizes drug resistant lung cancer cells to lysosomal pathway inhibitors. Theranostics 10, 2727–2743 (2020).

23. Paz, J. & Luders, J. Microtubule-Organizing Centers: Towards a Minimal Parts List. Trends Cell Biol 28, 176–187 (2018).

24. Bornens, M. Centrosome composition and microtubule anchoring mechanisms. Current opinion in cell biology 14, 25–34 (2002).

25. Azimzadeh, J. & Bornens, M. Structure and duplication of the centrosome. Journal of cell science 120, 2139–2142 (2007).

26. Kitagawa, D. et al. Spindle positioning in human cells relies on proper centriole formation and on the microcephaly proteins CPAP and STIL. J Cell Sci 124, 3884–3893 (2011).

27. Bettencourt-Dias, M., Hildebrandt, F., Pellman, D., Woods, G. & Godinho, S.A. Centrosomes and cilia in human disease. Trends Genet 27, 307–315 (2011).

28. Schmidt, T.I. et al. Control of centriole length by CPAP and CP110. Curr Biol 19, 1005–1011 (2009).

29. Tang, N. & Marshall, W.F. Centrosome positioning in vertebrate development. Journal of cell science 125, 4951–4961 (2012).

30. Pearson, C.G., Giddings, T.H., Jr. & Winey, M. Basal body components exhibit differential protein dynamics during nascent basal body assembly. Molecular biology of the cell 20, 904–914 (2009).

31. Kobayashi, T. & Dynlacht, B.D. Regulating the transition from centriole to basal body. J Cell Biol 193, 435–444 (2011).

32. Masyuk, T.V. et al. Centrosomal abnormalities characterize human and rodent cystic cholangiocytes and are associated with Cdc25A overexpression. The American journal of pathology 184, 110–121 (2014).

33. Thornton, G.K. & Woods, C.G. Primary microcephaly: do all roads lead to Rome? Trends Genet 25, 501–510 (2009).

34. Pelletier, L. & Yamashita, Y.M. Centrosome asymmetry and inheritance during animal development. Current opinion in cell biology 24, 541–546 (2012).

35. Hinchcliffe, E.H. & Sluder, G. "It takes two to tango": understanding how centrosome duplication is regulated throughout the cell cycle. Genes & development 15, 1167–1181 (2001).

36. Tanos, B.E. et al. Centriole distal appendages promote membrane docking, leading to cilia initiation. Genes & development 27, 163–168 (2013).

37. Ansari, D., Del Pino Bellido, C., Bauden, M. & Andersson, R. Centrosomal Abnormalities in Pancreatic Cancer: Molecular Mechanisms and Clinical Implications. Anticancer Res 38, 1241–1245 (2018).

38. Marteil, G. et al. Over-elongation of centrioles in cancer promotes centriole amplification and chromosome missegregation. Nat Commun 9, 1258 (2018).

39. Lopes, C.A.M. et al. Centrosome amplification arises before neoplasia and increases upon p53 loss in tumorigenesis. J Cell Biol 217, 2353–2363 (2018).

40. Faccion, R.S. et al. p53 expression and subcellular survivin localization improve the diagnosis and prognosis of patients with diffuse astrocytic tumors. Cell Oncol (Dordr) 41, 141–157 (2018).

41. Gudi, R., Haycraft, C.J., Bell, P.D., Li, Z. & Vasu, C. Centrobin-mediated regulation of the centrosomal protein 4.1-associated protein (CPAP) level limits centriole length during elongation stage. J Biol Chem 290, 6890–6902 (2015).

42. Gudi, R., Zou, C., Dhar, J., Gao, Q. & Vasu, C. Centrobin-centrosomal protein 4.1-associated protein (CPAP) interaction promotes CPAP localization to the centrioles during centriole duplication. The Journal of biological chemistry 289, 15166–15178 (2014).

43. Gudi, R., Zou, C., Li, J. & Gao, Q. Centrobin-tubulin interaction is required for centriole elongation and stability. J Cell Biol 193, 711–725 (2011).

44. Lee, M. & Rhee, K. Determination of Mother Centriole Maturation in CPAP-Depleted Cells Using the Ninein Antibody. Endocrinol Metab (Seoul) 30, 53–57 (2015).

45. Tang, C.J., Fu, R.H., Wu, K.S., Hsu, W.B. & Tang, T.K. CPAP is a cell-cycle regulated protein that controls centriole length. Nat Cell Biol 11, 825–831 (2009).

46. Januschke, J. et al. Centrobin controls mother-daughter centriole asymmetry in Drosophila neuroblasts. Nat Cell Biol 15, 241–248 (2013).

47. Song, L. et al. Inhibition of centriole duplication by centrobin depletion leads to p38-p53 mediated cell-cycle arrest. Cell Signal 22, 857–864 (2010).

48. Zou, C. et al. Centrobin: a novel daughter centriole-associated protein that is required for centriole duplication. J Cell Biol 171, 437–445 (2005).

49. Zheng, X. et al. Molecular basis for CPAP-tubulin interaction in controlling centriolar and ciliary length. Nat Commun 7, 11874 (2016).

50. Gabriel, E. et al. CPAP promotes timely cilium disassembly to maintain neural progenitor pool. EMBO J 35, 803–819 (2016).

51. Hehnly, H., Chen, C.T., Powers, C.M., Liu, H.L. & Doxsey, S. The centrosome regulates the Rab11-dependent recycling endosome pathway at appendages of the mother centriole. Curr Biol 22, 1944–1950 (2012).

52. Kohlmaier, G. et al. Overly long centrioles and defective cell division upon excess of the SAS-4-related protein CPAP. Curr Biol 19, 1012–1018 (2009).

53. Lee, M., Chang, J., Chang, S., Lee, K.S. & Rhee, K. Asymmetric spindle pole formation in CPAP-depleted mitotic cells. Biochem Biophys Res Commun 444, 644–650 (2014).

54. Cho, J.H., Chang, C.J., Chen, C.Y. & Tang, T.K. Depletion of CPAP by RNAi disrupts centrosome integrity and induces multipolar spindles. Biochem Biophys Res Commun 339, 742–747 (2006).

55. Wu, K.S. & Tang, T.K. CPAP is required for cilia formation in neuronal cells. Biol Open 1, 559–565 (2012).

56. Garcez, P.P. et al. Cenpj/CPAP regulates progenitor divisions and neuronal migration in the cerebral cortex downstream of Ascl1. Nat Commun 6, 6474 (2015).

57. McIntyre, R.E. et al. Disruption of mouse Cenpj, a regulator of centriole biogenesis, phenocopies Seckel syndrome. PLoS Genet 8, e1003022 (2012).

58. Al-Dosari, M.S., Shaheen, R., Colak, D. & Alkuraya, F.S. Novel CENPJ mutation causes Seckel syndrome. J Med Genet 47, 411–414 (2010).

59. Bond, J. et al. A centrosomal mechanism involving CDK5RAP2 and CENPJ controls brain size. Nat Genet 37, 353–355 (2005).

60. Gul, A. et al. A novel deletion mutation in CENPJ gene in a Pakistani family with autosomal recessive primary microcephaly. J Hum Genet 51, 760–764 (2006).

61. Sharma, A. et al. Centriolar CPAP/SAS-4 Imparts Slow Processive Microtubule Growth. Dev Cell 37, 362–376 (2016).

62. Mariappan, A. et al. Inhibition of CPAP-tubulin interaction prevents proliferation of centrosome-amplified cancer cells. EMBO J 38 (2019).

63. Xu, Q. et al. EGF induces epithelial-mesenchymal transition and cancer stem-like cell properties in human oral cancer cells via promoting Warburg effect. Oncotarget 8, 9557–9571 (2017).

64. Sato, F. et al. EGFR inhibitors prevent induction of cancer stem-like cells in esophageal squamous cell carcinoma by suppressing epithelial-mesenchymal transition. Cancer Biol Ther 16, 933–940 (2015).

65. Claperon, A. et al. EGF/EGFR axis contributes to the progression of cholangiocarcinoma through the induction of an epithelial-mesenchymal transition. J Hepatol 61, 325–332 (2014).

66. Sabat-Pospiech, D., Fabian-Kolpanowicz, K., Prior, I.A., Coulson, J.M. & Fielding, A.B. Targeting centrosome amplification, an Achilles’ heel of cancer. Biochem Soc Trans 47, 1209–1222 (2019).

67. Pihan, G.A. Centrosome dysfunction contributes to chromosome instability chromoanagenesis, and genome reprograming in cancer. Front Oncol 3, 277 (2013).

68. Fan, G. et al. Loss of KLF14 triggers centrosome amplification and tumorigenesis. Nat Commun 6, 8450 (2015).

69. Zhang, Z., Dong, Z., Lauxen, I.S., Filho, M.S. & Nor, J.E. Endothelial cell-secreted EGF induces epithelial to mesenchymal transition and endows head and neck cancer cells with stem-like phenotype. Cancer Res 74, 2869–2881 (2014).

70. Thiery, J.P. Epithelial-mesenchymal transitions in tumour progression. Nat Rev Cancer 2, 442–454 (2002).

71. Kang, Y. & Massague, J. Epithelial-mesenchymal transitions: twist in development and metastasis. Cell 118, 277–279 (2004).

72. Muthusami, S., Prabakaran, D.S., Yu, J.R. & Park, W.Y. EGF-induced expression of Fused Toes Homolog (FTS) facilitates epithelial-mesenchymal transition and promotes cell migration in ME180 cervical cancer cells. Cancer Lett 351, 252–259 (2014).

73. Miettinen, P.J., Ebner, R., Lopez, A.R. & Derynck, R. TGF-beta induced transdifferentiation of mammary epithelial cells to mesenchymal cells: involvement of type I receptors. J Cell Biol 127, 2021–2036 (1994).

74. Hung, L.Y., Tang, C.J. & Tang, T.K. Protein 4.1 R-135 interacts with a novel centrosomal protein (CPAP) which is associated with the gamma-tubulin complex. Mol Cell Biol 20, 7813–7825 (2000).

75. Hung, L.Y., Chen, H.L., Chang, C.W., Li, B.R. & Tang, T.K. Identification of a novel microtubule-destabilizing motif in CPAP that binds to tubulin heterodimers and inhibits microtubule assembly. Molecular biology of the cell 15, 2697–2706 (2004).

76. Costa, V. et al. EGFR amplification and expression in oral squamous cell carcinoma in young adults. Int J Oral Maxillofac Surg 47, 817–823 (2018).

77. Taoudi Benchekroun, M. et al. Epidermal growth factor receptor expression and gene copy number in the risk of oral cancer. Cancer Prev Res (Phila) 3, 800–809 (2010).

78. Guo, M. et al. Comparison of the expression of TGF-beta1, E-cadherin, N-cadherin, TP53, RB1CC1 and HIF-1alpha in oral squamous cell carcinoma and lymph node metastases of humans and mice. Oncol Lett 15, 1639–1645 (2018).

79. Chen, M.F., Wang, W.H., Lin, P.Y., Lee, K.D. & Chen, W.C. Significance of the TGF-beta1/IL-6 axis in oral cancer. Clin Sci (Lond) 122, 459–472 (2012).

80. Peltanova, B., Raudenska, M. & Masarik, M. Effect of tumor microenvironment on pathogenesis of the head and neck squamous cell carcinoma: a systematic review. Mol Cancer 18, 63 (2019).

81. Ang, K.K. et al. Impact of epidermal growth factor receptor expression on survival and pattern of relapse in patients with advanced head and neck carcinoma. Cancer Res 62, 7350–7356 (2002).

82. Oliveira, S., van Bergen en Henegouwen, P.M., Storm, G. & Schiffelers, R.M. Molecular biology of epidermal growth factor receptor inhibition for cancer therapy. Expert Opin Biol Ther 6, 605–617 (2006).

83. Thariat, J. et al. Epidermal growth factor receptor protein detection in head and neck cancer patients: a many-faceted picture. Clin Cancer Res 18, 1313–1322 (2012).

84. Egloff, A.M. & Grandis, J.R. Targeting epidermal growth factor receptor and SRC pathways in head and neck cancer. Semin Oncol 35, 286–297 (2008).

85. Mahipal, A., Kothari, N. & Gupta, S. Epidermal growth factor receptor inhibitors: coming of age. Cancer Control 21, 74–79 (2014).

86. Loeffler-Ragg, J., Schwentner, I., Sprinzl, G.M. & Zwierzina, H. EGFR inhibition as a therapy for head and neck squamous cell carcinoma. Expert Opin Investig Drugs 17, 1517–1531 (2008).

87. Birchmeier, W. & Birchmeier, C. Epithelial-mesenchymal transitions in development and tumor progression. EXS 74, 1–15 (1995).

88. Kimura, I. et al. Loss of epidermal growth factor receptor expression in oral squamous cell carcinoma is associated with invasiveness and epithelial-mesenchymal transition. Oncol Lett 11, 201–207 (2016).

