## Supplemental data for "Loss of CPAP expression promotes sustained EGFR signaling and Epithelial-Mesenchymal Transition in oral cancer cells"

**Supplemental Fig. 1**

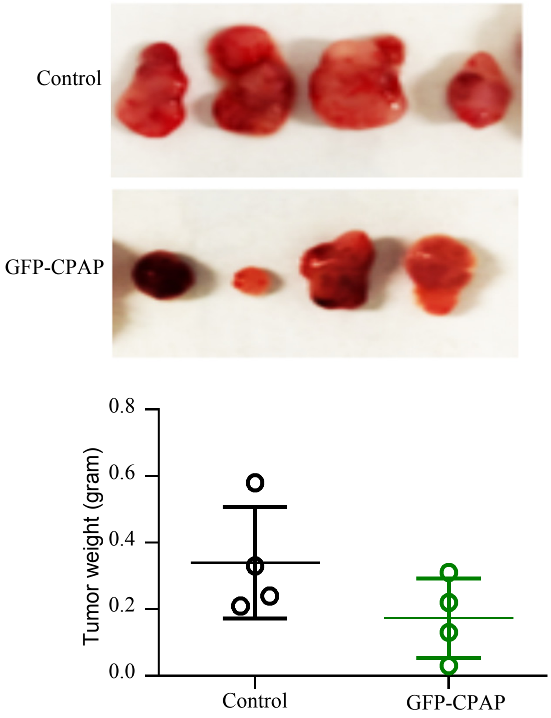


**Supplemental Fig. 1: CPAP overexpression enhances tumorigenic property of OSCC cells.** UM-SCC-74B cells stably expressing control vector or GFP-CPAP (under doxycycline inducible promoter) vector were subcutaneously injected in nude mice (4 mice/group). Day 24 post-injection, tumors were harvested and weighed.
